# Behavioral and genetic markers of susceptibility to escalate fentanyl intake

**DOI:** 10.1101/2024.12.06.627259

**Authors:** Jack Keady, Richard Charnigo, Jakob D. Shaykin, Emily R. Prantzalos, Mengfan Xia, Emily Denehy, Cody Bumgardner, Justin Miller, Pavel Ortinski, Michael T. Bardo, Jill R. Turner

## Abstract

**Background:** The “loss of control” over drug consumption, present in opioid use disorder (OUD) and known as escalation of intake, is well-established in preclinical rodent models. However, little is known about how antecedent behavioral characteristics, such as valuation of hedonic reinforcers prior to drug use, may impact the trajectory of fentanyl intake over time. Moreover, it is unclear if distinct escalation phenotypes may be driven by genetic markers predictive of OUD susceptibility.

**Methods:** Male and female Sprague-Dawley rats (n=63) were trained in a sucrose reinforcement task using a progressive ratio schedule. Individual differences in responsivity to sucrose were hypothesized to predict escalation of fentanyl intake. Rats underwent daily 1-h acquisition sessions for i.v. fentanyl self-administration (2.5 µg/kg; FR1) for 7 days, followed by 21 6-h escalation sessions, then tissue from prefrontal cortex was collected for RNA sequencing and qPCR. Latent growth curve and group-based trajectory modeling were used, respectively, to evaluate the association between sucrose reinforcement and fentanyl self-administration and to identify whether distinct escalation phenotypes can be linked to gene expression patterns.

**Results:** Sucrose breakpoints were not predictive of fentanyl acquisition nor change during escalation, but did predict fentanyl intake on the first day of extended access to fentanyl. Permutation analyses did not identify associations between behavior and single gene expression when evaluated overall, or between our ascertained phenotypes. However, weighted genome correlation network analysis (WGCNA) and gene set enrichment analysis (GSEA) determined several gene modules linked to escalated fentanyl intake, including genes coding for voltage-gated potassium channels, calcium channels, and genes involved in excitatory synaptic signaling. Transcription factor analyses identified EZH2 and JARID2 as potential transcriptional regulators associated with escalated fentanyl intake. Genome-wide association study (GWAS) term categories were also generated and positively associated with terms relating to substance use disorders.

**Discussion:** Escalation of opioid intake is largely distinct from motivation for natural reward, such as sucrose. Further, the gene networks associated with fentanyl escalation suggest that engagement of select molecular pathways distinguish individuals with “addiction prone” behavioral endophenotypes, potentially representing druggable targets for opioid use disorder. Our extended in silico identification of SNPs and transcription factors associated with the “addiction prone” high escalating rats highlights the importance of integrating findings from translational preclinical models. Through a precision medicine approach, our results may aid in the development of patient-centered treatment options for those with OUD.

## INTRODUCTION

Escalation of intake is a prominent feature of opioid use in humans. Diagnostic criteria of opioid use disorder (OUD) incorporate measures of increased drug consumption, such as changes in frequency or dose, as a factor directly impacting severity of opioid use across time. While increased opioid use may be perceived as tolerance to a unit dose of the drug, clinical data indicate that the small changes in pharmacokinetic parameters for orally administered opioids after chronic use (*1, 2*) are insufficient to explain the 10-20 fold increase of drug intake common with escalation (*3*). In the meantime, escalated intake of opioids has been linked to a significantly elevated risk of developing both opioid and non-opioid substance use disorders, as well as other adverse outcomes, in a recent retrospective study of over 50,000 chronic pain patients in the Veterans Healthcare Administration system (*4*). These considerations suggest that escalation of opioid use represents a phenotypic signature that can predict substance use in the future and may potentially capture other traits characteristic of risk.

Escalation of drug intake can be observed in rodent models under conditions of extended access to opioids in self-administration (SA) sessions. Extended access SA sessions last anywhere from 3-23 h, similar to the time window of opioid use in people with OUD (*5*), whereas short (1-2 h) SA sessions serve as a useful assay for controlled, but limited intake (*6, 7*). Moreover, extended access SA sessions are advantageous in their ability to produce increased heroin-and fentanyl-induced reinstatement and heightened signs of withdrawal compared to rodents with short access (*8–10*). Many factors may influence escalated use, including intermittency of access, hedonic value changes, withdrawal symptoms, habit formation, and environmental (e.g., context or social) events (*11*). While contributing to significant heterogeneity in behavioral patterns associated with fentanyl use, these observations also highlight the possibility that behavioral characteristics present prior to initiation of fentanyl use may impact the emergence of OUD.

The prefrontal cortex (PFC) is an important site of both neuronal activity underlying addictive behavior and changes in gene expression induced by opioid exposure (*12–14*). Widespread transcriptional activation in the PFC serves as a regulatory response following chronic opioid exposure (*15*). Moreover, while acute opioid exposure has acute pharmacological actions throughout the central nervous system (CNS), counter-adaptational transcriptomic signatures persist in the PFC (*16*). Therefore, analyses of PFC genomic changes provide a lens through which individual fentanyl escalation phenotypes may be captured, with the goal of uncovering the neural mechanisms underlying the “loss of control” phenomenon.

In the present study, we aimed to: 1) examine whether individual differences in motivation for non-drug reward influence the likelihood and extent of subsequent fentanyl use, 2) characterize individual differences in fentanyl escalation, and 3) determine the transcriptomic signatures associated with escalation-prone behavioral phenotypes.

## METHODS

### Subjects

Male and female Sprague-Dawley rats (PND 50-55; n=72; n=36 male; n=36 female) were individually housed in a colony room with *ad libitum* water access and were maintained on a 12-h/12-h light/dark cycle with lights on at 0700 hours. All experimental procedures were followed in accordance with the NIH Guide for the Care and Use of Laboratory Animals (8^th^ edition, 2011) and approved by the University of Kentucky IACUC.

### Sucrose self-administration

Self-administration experiments were conducted in ventilated, sound attenuating operant chambers, equipped with a house light, two nose-poke holes, a pellet dispenser and a food receptacle (Med-Associates Inc., East Fairfield, VT, USA). During the experiment (**Figure 1.A**), rats had *ad libitum* access to water and were given 20 g of normal chow per day after each operant session. All subjects were trained daily on a fixed-ratio one (FR1) schedule of reinforcement, in which each nose poke into a reinforced active hole delivered a single 45 mg sucrose pellet. Pokes into the inactive hole had no programmed consequences. Each sucrose pellet was followed by a 20 s timeout period during which house light went off and nose pokes had no scheduled consequences. After 10 days of FR1 nose-poke training, the rats progressed to the FR3 schedule (three nose pokes for one sucrose pellet) for 3 days and then to FR10 (ten nose pokes for one sucrose pellet) for 8 days. Following FR training, the rats were placed on a progressive ratio (PR) schedule of reinforcement during which successive sucrose pellets could be earned according to an increasing number of reinforced nose-pokes based on the formula: [5e^(pellet^ ^#^ * ^0.2)^] – 5 (*17*). The session ended when rats failed to reach the next nose-poke criterion within 1 h. The final reinforced nose-poke criterion achieved was recorded as the “breakpoint” value for that session. On the last day of sucrose PR, the rats were given *ad libitum* chow.

**Figure 1.**
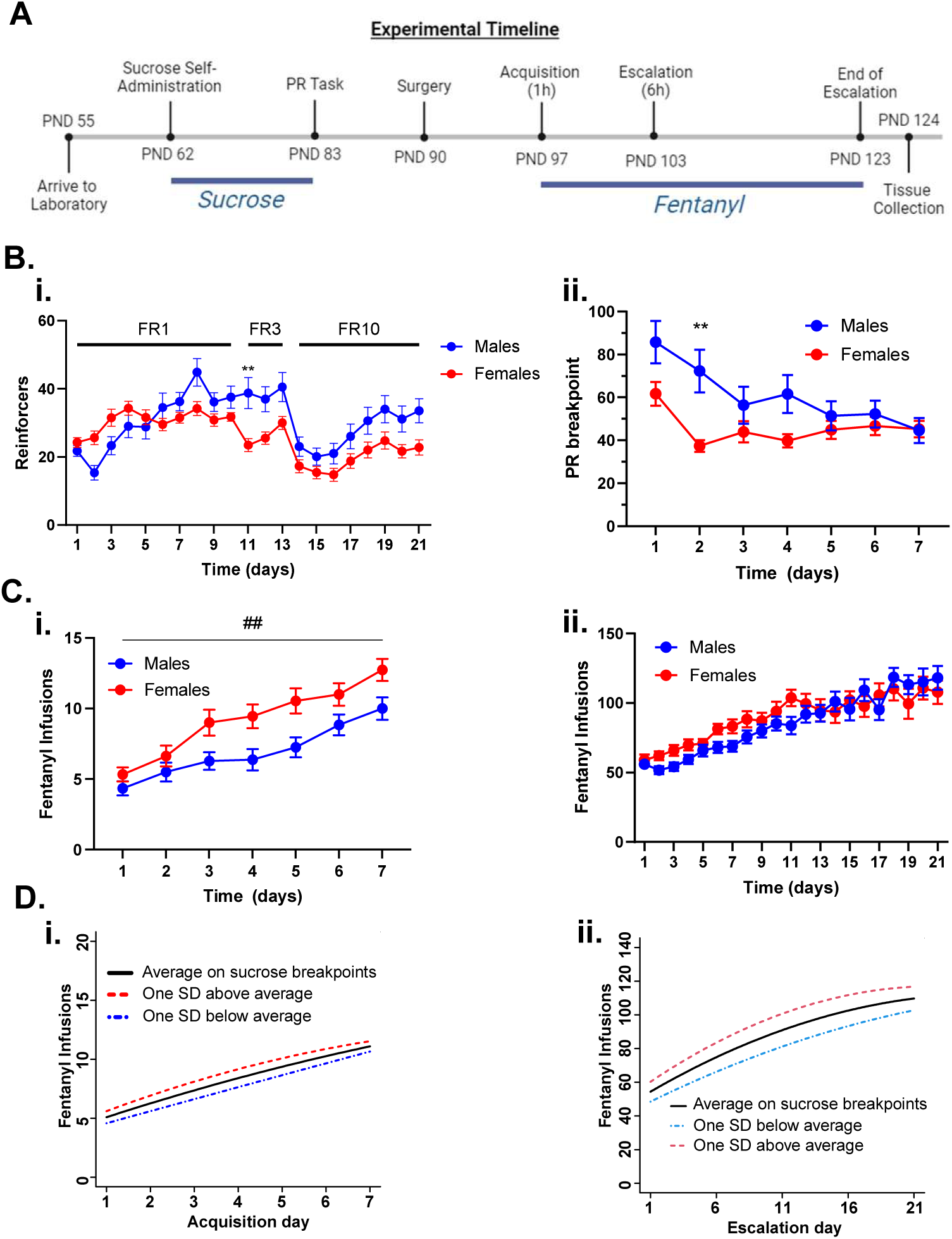
Motivation for Sucrose as Predictor of Fentanyl Escalation. (**A**) Experimental timeline. (**B**) Sucrose self-administration. i) PND (62–82) Number of sucrose pellets that male (blue circles) and female (red circles) rats received over the course of FR 1 (day 1-10), FR 3 (day 11-13), and FR 10 (day 14-21) sessions. Ii) Progressive ratio breakpoints for male and female rats across the 7 days of PR training. (**C**) Fentanyl self-administration. Rats underwent i) 1 h acquisition and ii) 6 h escalation sessions for fentanyl (2.5 µg/kg/inf). (**D**) Latent growth curve model considering hypothetical rats that behave at exactly the average breakpoint (black line) over the last three days of sucrose PR, at one standard deviation above the average (red line), and at one standard deviation below the average (blue line). The relationship between sucrose PR and fentanyl infusions is visually appreciated as the magnitude of difference between the 3 growth curves [ ** indicates p < 0.01 as determined by Bonferroni’s multiple comparison tests, ^##^ indicates p < 0.01 for the main effect of sex. Data in figures 1B and 1C represent mean±SEM].

### Catheterization and Fentanyl Self-Administration

On PND 90, rats were anesthetized with a ketamine/xylazine/acepromazine (75, 7.5, 0.75 mg/kg) mixture at a 0.15 ml/100g body weight (i.p.) and were implanted with chronic indwelling jugular catheters with a cannula secured to the skull with dental acrylic anchored by four screws. Rats were given 5-7 days to recover after surgery and catheters were flushed daily with a solution containing gentamycin, heparin, and saline. Rats were then trained to self-administer intravenous (i.v.) fentanyl (2.5 µg/kg/infusion) using a two-lever operant procedure under FR 1 schedule of reinforcement in a two-lever operant conditioning chamber (ENV-001; MED Associates, St. Albans, VT) using methods described previously by Malone *et al*. (2021)(*18*). Briefly, young adult male and female Sprague-Dawley rats (55-60 days of age) were trained to self-administer fentanyl (2.5 µg/kg/infusion) in a 2-lever operant conditioning chamber using an autoshaping procedure. During the first 35 min of a 1-hr session, one lever (active lever) was extended into the chamber, followed 15 sec later by retracting the lever and delivering an infusion accompanied by 20-sec illumination of 2 cue lights, one located above each lever; pressing the lever during the 15-sec extension led to immediate delivery of the infusion and cue illumination. The intention of this procedure was to train rats to approach the active lever, while controlling for the number of drug exposures across rats. Two hr following the autoshaping session, rats were placed back into the chamber with both levers continuously extended for 1-hr and allowed to earn a fentanyl infusion and 20-sec cue light illumination with each active lever press (FR1); pressing on the active lever during the 20-sec cue light illumination was not reinforced. This protocol occurred across 7 consecutive days, at which point autoshaping sessions were omitted and the FR1 session was extended to 6 hr across 21 days.

### Behavioral Data Analysis

*General remarks*. Data analysis, visualization, and statistical calculations were carried out using SAS (Version 9.4), R (Version 3.4.1 or later), GraphPad Prism (version 9.4 or later) and MATLAB (R2022A); the PROC TRAJ add-on to SAS (trajv9-64) (*19*) was employed, as was the scatterplot3D package for R (version 0.3-44) (*20*). Formal hypothesis tests used alpha = 0.05. Further details about methods appear in the supplemental material.

*Quantifying sucrose preference*. Breakpoint values were recorded for each rat throughout the 7-day sucrose PR protocol. The average breakpoint over the last three days was calculated for each rat, and these three-day averages were converted to *z* scores by subtraction of the overall mean (47.5) and then division by the overall standard deviation (29.8).

*Latent growth curve modeling of fentanyl acquisition and escalation*. We used PROC MIXED in SAS (Version 9.4) to fit a latent growth curve model to describe the expected number of fentanyl infusions as a quadratic function of time, with coefficients of the quadratic function (hereafter, “coefficients”) permitted to depend linearly on sucrose PR protocol *z-scores*. Fentanyl acquisition and escalation phases were modeled separately. Partial data were included from rats not having valid information for all 7 days (acquisition phase) or all 21 days (escalation phase). The latent growth curve model included random effects for rat-specific adjustments to the coefficients with an unstructured covariance matrix.

*Group-based trajectory modeling (escalation phase)*. We used the PROC TRAJ add-on to SAS (*20*) with the MULTGROUPS option to perform group-based trajectory modeling simultaneously for three outcomes measured daily during the 21-day escalation phase: active lever presses resulting in fentanyl infusions (reinforced presses), nonreinforced active lever presses during signaled timeouts (futile presses), and non-reinforced inactive lever presses (inactive presses). Partial data were included from rats not having valid information for all 21 days. Futile presses and inactive presses were sufficiently non-normal in their distributions that we log transformed them (after the addition of 1, to avoid taking a logarithm of 0) before applying PROC TRAJ under censored normal model specifications (with upper limits of 500 for reinforced presses and 10 for the log transformed outcomes); however, for ease of interpretation, figures in the main manuscript and supplemental material convey the original, non-transformed values.

*Group-based trajectory modeling (acquisition phase)*. The group-based trajectory modeling methodology was similar to that for the escalation phase, with the following exceptions. First, because non-normality was less pronounced in the acquisition phase data, the futile press and inactive press outcomes were not log transformed. Second, the censored normal model specifications were altered (with upper limits of 30 for reinforced presses, 70 for futile presses, and 40 for inactive presses). Third, we adopted a 5 group model. For further details on the classification of the five groups, please refer to the “Group-Based Trajectory Modeling” section in the Results. Fourth, the backward elimination procedure (**Suppl. File 1**) resulted in only three of the quadratic trajectories being reduced to linear.

*Cross-classifications into trajectory groups*. We used PROC FREQ in Version 9.4 of SAS with the EXACT option to test for association between trajectory group assignments derived from the escalation and the acquisition experiments; this test was based on the 56 rats assigned by trajectory modeling to non-anomalous (e.g N=1) groups in both experiments.

### Tissue Collection, RNA-Sequencing, Secondary Analysis

*Tissue Collection and RNA Extraction.* Eighteen h after the last self-administration session, rats were euthanized, and brain tissue was collected. Total RNA was extracted from the PFC using TRIzol (Invitrogen) and an RNeazy Mini Kit (QIAGEN) according to the manufacturers’ instructions. RNA samples were quantified using the NanoDropOne (Thermo Scientific) and were confirmed to be of good quality (RNA 260/280 >2.0).

*RNA-Sequencing and Alignment.* Tissue from rats classified into escalation Groups 1 (M = 5, F =3), 2 (M=4, F =4), and 3 (M = 4, F =4) were selected for RNA-sequencing. These 24 samples were sent to GENEWIZ from Azenta Life Sciences (South Plainfield, NJ) for next-generation sequencing (standard RNA-seq service), The remaining samples were used as biological replicates for validation via RT-PCR: N = 4 - 7 per escalation group (Group 1: N =4 [M=1,F=3]; Group 2: N=7 [M=6, F =1]; Group 3: N=5 [M=1, F=4]). Sample quality was verified (RNA integrity numbers >9) and RNA libraries were prepped with PolyA selection before sequencing using an Illumina HiSeq platform with 2 × 150 base pair (bp) paired-end reads. RNA - sequencing data was preprocessed and analyzed for differential expression using Galaxy (24.0.1.dev0), a web-based cloud computing platform for bioinformatic analysis, as previously described.(*21–24*) Briefly, fastq files from the Illumina Hiseq4000 were checked for quality using FastQC (Galaxy Version 0.74+galaxy0). Trim Galore (Galaxy Version 0.6.7+galaxy0) was used to remove adaptor content, low quality reads (<20 PHRED score), and read length (<20 bp) from fastq files, between 0.3% to 0.4% of reads were removed from each sample for not meeting quality requirements. Trim Galore output was aligned to the rat genome (rn6, downloaded from iGenome, Illumina) using STAR (Galaxy Version 2.7.11a+galaxy0). Average input read counts for the rat PFC samples were 29.39 million, with an average of 80.85% uniquely mapped reads. Gene counts were determined using featureCounts (Galaxy Version 2.0.3+galaxy2).

*Differential Expression*. Differential RNA expression was determined using pyDESeq2 (0.4.9) at a false discovery rate (FDR) < 0.05, filtering out genes with less than 10 total reads per row, with fentanyl escalation groups treated as the between-group comparisons (*25*). Sex was included as a variable in differential expression, but not treated as a between-group factor due to lack of sex differences observed behaviorally. Information regarding permutation-based testing for single gene differential expression can be found in **Suppl. File 1**.

*Weighted Gene Co-Expression Network Analysis*. We used the Python package PyWGCNA (2.0.3) to conduct Weighted Gene Co-Expression Network Analysis (WGCNA) on the sequencing results with minimum module size = 30, MEDissThres = 0.2, and a soft power 17 (**Suppl. Fig. 1**)(*26*) on the corrected counts from pyDESeq2’s median of ratio. Functional enrichment of modules identified via WGCNA were performed via the GSEAPY (1.1.3) enrichr function (*27, 28*), using the 2023 Gene Ontologies (Biological Process, Cellular Component, and Molecular Function)(*29, 30*) as well as the 2023 GWAS catalog (*31*) with all gene names within a given module as input.

*Transcription Factor Analysis*. Binding Analysis for Regulation of Transcription (BART2 v=2.1) was used to predict potential transcription factors regulating each module, using published mouse and human chromatin immunoprecipitation sequencing libraries (*32*).

### Quantitative RT-PCR

Quantitative reverse transcriptase PCR was performed as described previously (*33–35*) on PFC samples from escalation Groups 1,2, and 3. Briefly, RNA was isolated using the RNeasy Mini kit (Qiagen) and complementary DNA (cDNA) was synthesized from 500 ng of isolates RNA. qPCR reactions were assembled using Thermo Scientific Maxima SYBR Green master mix, cDNA, and primers diluted down to final concentration of 5nM (Integrated DNA Technologies, Coralville, IA) designed via Primer3Plus. Levels of mRNA were determined using the 2^-ΔΔCT^ method (*36*), and genes of interest were normalized to the housekeeping gene TATA-Binding Protein (*tbp*) after assessing the genes of interest at values in relation to *tbp* and Glyceraldehyde 3-phosphate dehydrogenase (*gapdh*). Gene expression values were normalized to fentanyl escalating Group 1, and primer sequences are shown in **Table 1**. Genes chosen for RT-PCR validation were selected by generating a priority score by combining results from the differential expression analysis and WGCNA with the following formula:

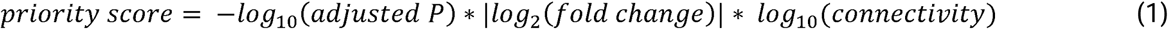

**Table 1.**
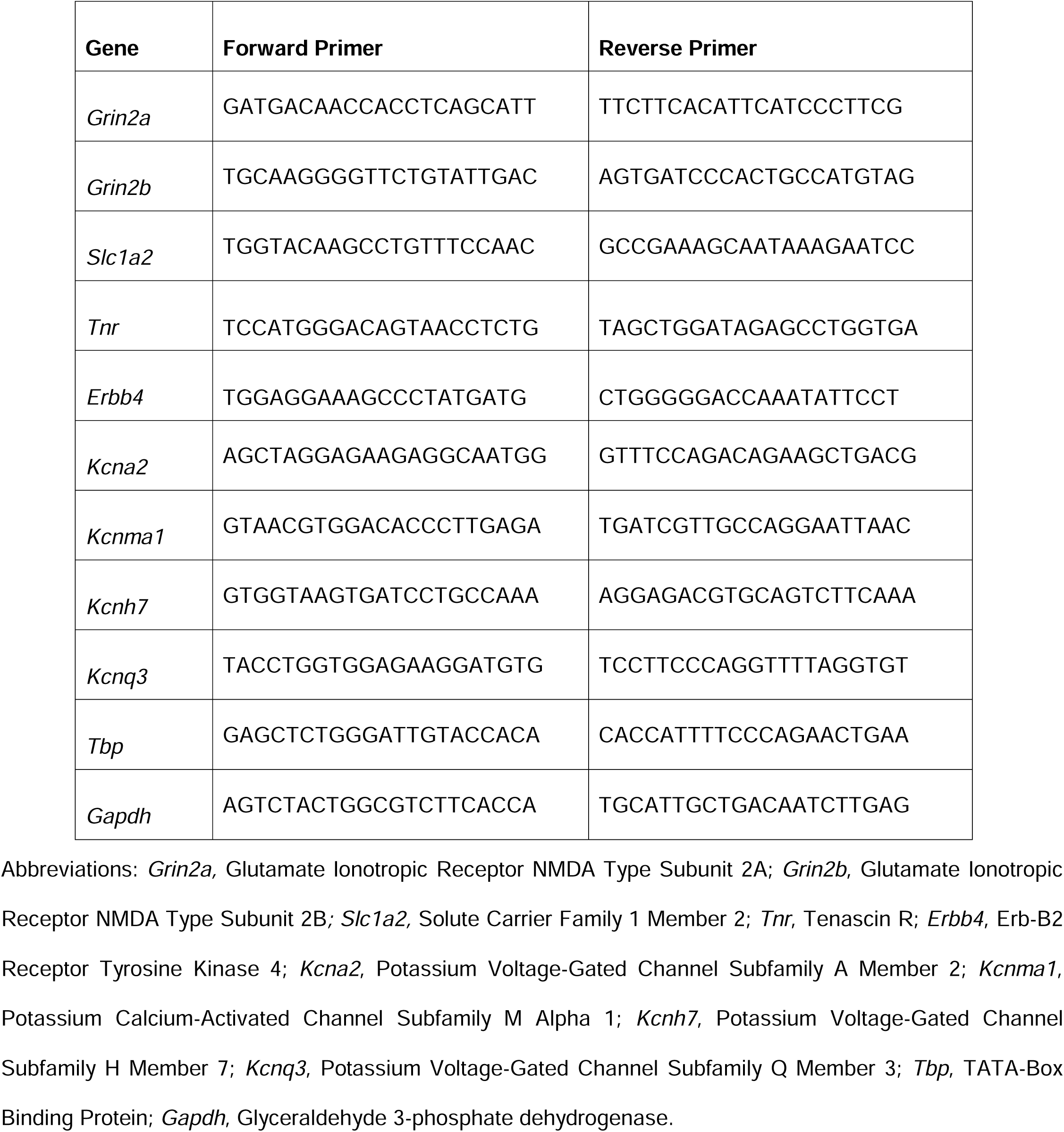
Sequence of primers used in qPCR.

Where *adjusted P* is the adjusted P value and |*log_2_(fold change*)| is the absolute log2(FC) for a given gene between two escalation phenotypes, and connectivity is the sum of connection strength of a given gene with all other genes within the same module generated by the WGCNA package (*26, 37*). Priority scores were generated for genes in modules that correlate to our observed fentanyl escalation phenotypes. This formula was chosen to generate priority scores as it allows us to prioritize genes for RT-PCR validation that have differential expression, or trends towards differential expression, between observed phenotypes and are highly connected within a module correlated to our observed phenotypes. Therefore, the priority score reflects genes with potential role in mediating shifts in the transcriptomic profiles of the escalation phenotype.

Differences in gene expression between escalation groups were tested using a one-way ANOVA in GraphPad Prism 10.0 software package (GraphPad Software, CA). RT-PCR data were normalized to escalation group 1 (non-escalator). To examine relationship between qPCR results and escalation phenotypes, a combined *z-score* for the biological replicates including reinforcers, inactive responses, and futile responding for each rat was calculated over the last 7 self-administration sessions using the following formula:

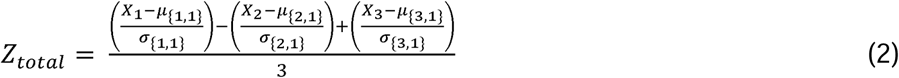

Where *X*_1_ is reinforcers, X_2_ is futile presses, and X_3_ is inactive presses, all averaged for each individual rat across the last 7 self-administration sessions. Then let *µ*{*k*,1} and σ_{*k*,1}_ represent mean and standard deviation of the kth outcome (k =1,2,3) for group 1 biological replicates. SPSS Statistics software package 29 (IBM, NY) was then used to perform linear regressions treating qPCR data as the independent variables [9 genes of interest] and Ztotal as the dependent variable. The stepwise method with F value of 2 for entry and 1 for removal was used in these models.

We also performed a principal component analysis via scikit-learn (1.5.1) on the biological replicates, with *X*_1_, *X*_2_, and *X*_3_. SPSS Statistics software package 29 (IBM, NY) was then used to perform linear regressions treating qPCR data as the independent variables [9 genes of interest] and the first principal component of (PC1) as the dependent variable. The stepwise method with F value of 2 for entry and 1 for removal was used in these models.

Additional supplementary methods can be found in **Suppl. File 1**.

## RESULTS

### Sucrose and Fentanyl Self-Administration

We completed behavioral profiling of sucrose self-administration on a PR schedule of reinforcement in 72 rats (36 males, 36 females; **Suppl. Fig. 2)** that were tested in 6 squads of 12 rats over an 8-month period. **Figure 1.Bi** displays reinforcers earned across 21 days of sucrose self-administration, prior to the PR task. A mixed-effects ANOVA indicated a significant sex x time interaction (F (20, 1398) = 4.83, p < 0.001). To probe the interaction, a Bonferroni multiple comparisons test determined that male rats earned significantly more sucrose pellets on session 11 compared to females (p = 0.002). **Figure 1.Bii** displays sucrose breakpoints as a function of time. A mixed-effects ANOVA indicated a significant sex x time interaction (F (6, 420) = 7.091, p < 0.001), and a Bonferroni multiple comparisons test indicated a significant difference in PR breakpoints for sucrose between males and females at session 2 (p = 0.001). The data indicated a decrease of sucrose breakpoints from the first to the second and third (for males) day of PR training followed by a more gradual decrease for males only over the remaining days. Average breakpoint values across the last 3 days of PR training for each rat were used for associations with fentanyl seeking in subsequent analyses.

After seven days of sucrose PR, the rats went on to fentanyl self-administration training, progressing from 1-h sessions for 7 days (“acquisition phase”, **Fig. 1.Ci**) to 6-h sessions for 21 days (“escalation phase”, **Fig. 1.Cii**). A total of 63 rats (n=32 male; n=31 female) completed all self-administration sessions; rats excluded from these analyses were due to catheter patency or surgical failure. **Figure 1.Ci** displays fentanyl self-administration during the acquisition phase. A mixed-effects ANOVA indicated a significant main effect of sex (F (1, 63) = 7.70, p = 0.007) and time (F (6, 374) = 47.17, p < 0.001), such that intake increased across the 7 days for both sexes. Figure 1.Cii displays fentanyl self-administration during the escalation phase. A mixed-effects analysis indicated a significant effect of time (F (20, 952) = 22.68, p < 0.001), such that intake increased across the 21 days. Sex differences seen in acquisition were not observed in the escalation phase (F (1, 56) = 1.80, p = 0.19). Individual data points for reinforced, inactive, and futile presses can be found in **Suppl. Fig 3**.

### Latent growth curve modeling

To evaluate whether sucrose PR performance predicted fentanyl seeking, we used latent growth curve modeling. We quantified a pattern of reinforced lever pressing for fentanyl as the initial number of presses at day 1 of training as well as the change in number of presses through the subsequent training days, with both allowed to depend on sucrose preference. We found that sucrose preference did not predict the overall pattern of reinforced presses during acquisition (p = 0.53; **Fig. 1.Di**), but did predict the overall pattern of reinforced pressing during escalation (p = 0.04; **Fig. 1.Dii**). Interestingly, for the escalation phase, it was the initial number of reinforced presses (p = 0.03), rather than the change in reinforced pressing across sessions (p = 0.67), that was associated with antecedent sucrose PR performance. More specifically, latent growth curve modeling predicted 54 initial fentanyl infusions for a rat with average sucrose PR performance and a difference of ∼6 reinforced presses for each standard deviation increase in sucrose PR performance at day 1 of escalation. On the last day of escalation, each standard deviation increase in sucrose PR breakpoint predicted only a marginally larger difference of 7 presses, highlighting that the change in lever pressing over the escalation phase was not associated with antecedent sucrose PR performance.

We also pursued latent growth curve modeling to test for associations between sucrose PR and inactive or futile lever pressing during the acquisition and escalation phases of fentanyl self-administration. During the acquisition phase, sucrose PR did not predict inactive (p = 0.93) or futile (p = 0.84) lever pressing (**Suppl. Fig. 4A-B**). Likewise, during the escalation phase, sucrose PR did not predict inactive (p = 0.61) or futile (p = 0.53) lever pressing (**Suppl. Fig. 4C-D**). Thus, these results support a relationship specifically between sucrose PR and reinforced lever pressing resulting in fentanyl infusions during the first session of the escalation phase, but not between sucrose PR and futile presses on the active lever, nor between sucrose PR and inactive lever presses regardless of phase of self-administration training.

### Group-based trajectory modeling

We also investigated whether sucrose PR performance could predict membership in groups of phenotypically similar rats. To this end, we used group-based trajectory modeling to probabilistically cluster rats based on the combination of reinforced, futile, and inactive lever presses.

For the acquisition phase, group-based trajectory modeling identified five behavioral phenotypes, exhibited by n=15 (group 1), n=21 (group 2), n=6 (group 3), n=20 (group 4), and n=1 (group 5) rats respectively. Group 5 contained one rat and was excluded from further analysis over doubts that behavior of a single rat is generalizable, but lever press data from this group is nevertheless presented in **Suppl. Fig. 5.** The remaining group numbers reflect fentanyl intake in the ascending order (**Fig. 2A-C**). Rats in group 1 maintained relatively stable levels of reinforced and futile presses, whereas rats in group 4 reached a plateau of reinforced presses by day 5-6 of fentanyl acquisition with stable futile presses. Rats in groups 2 and 3 displayed near linear increases in reinforced lever pressing at similar levels, but differed on futile presses. While futile press numbers in group 2 were stable, futile presses in group 3 approximately tripled over the seven days of acquisition training (**Fig. 2C**). As for inactive presses, group 3 and group 2 began at high and low levels, respectively, with intermediate (and nearly identical) numbers of inactive presses by groups 1 and 4. All groups converged on similar numbers of inactive presses by the last day (**Fig. 2B**). We found no significant association between sucrose PR *z-scores* and acquisition group assignment (p = 0.43; **Fig. 2D**).

**Figure 2.**
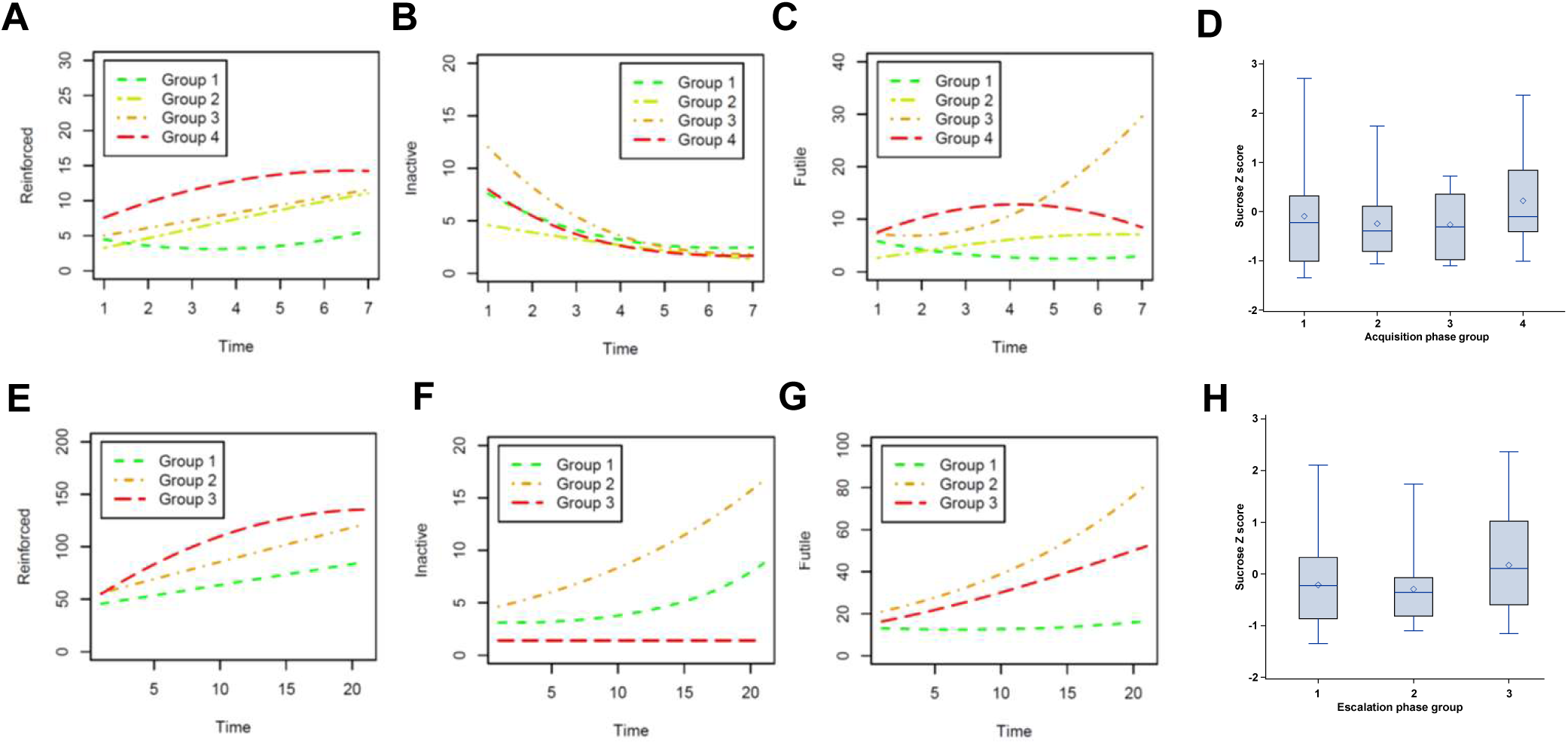
**Group based trajectory modeling**. Group assignment for the acquisition phase was based on the following parameters: (**A**) reinforced lever presses, (**B**) inactive lever presses, (**C**) futile lever presses for rats placed in Group 1 (green), Group 2 (yellow), Group 3 (orange), and Group 4 (red). A separate model demonstrating group assignment for the escalation phase based on (**E**) reinforced lever presses, (**F**) inactive lever presses, (**G**) futile lever presses for rats placed in Group 1 (green), Group 2 (orange), Group 3 (red). Panels (**D**) and (**H**) show box plots of sucrose breakpoint *z-scores* by assigned acquisition and escalation groups.

For the escalation phase, group-based trajectory modeling discovered four behavioral phenotypes, expressed by n=17 (group 1), n=24 (group 2), n=15 (group 3), and n=2 (group 4) rats, respectively. Again, because of dubious generalizability, group 4 was largely excluded from further analysis (**Fig. 2E-G**). The remaining group numbers were assigned to represent increasing fentanyl intake, such that escalation groups 1, 2, and 3 can be characterized as low, medium, and high escalators, respectively. Reinforced presses increased nearly linearly for low and medium escalators but plateaued in high escalators by day 16-17 (**Fig. 2E**). Medium and high escalators, but not low escalators, increased futile lever pressing (**Fig. 2G**). Whereas high escalators maintained a low level of inactive presses, low and medium escalators increased their inactive lever pressing (**Fig. 2F**). Assignment to escalation groups was not clearly associated with sucrose PR (p = 0.18; **Fig. 2H**). However, the excluded group 4 rats had the highest sucrose PR performance (**Suppl. Fig. 6A**), had extreme numbers of futile presses and displayed parabolic patterns of reinforced, inactive, and futile presses (**Suppl. Fig. 6B-D)**. Three-dimensional scatterplots to show acquisition and escalation trajectories of reinforced, futile, and inactive pressing, as well as animations to depict these trajectories from different perspectives, can be found in **Suppl. Fig. 7**.

Finally, we examined whether sex was associated with group assignments generated by group-based trajectory modeling. Acquisition group membership was significantly associated with sex (p=0.001), such that low fentanyl intake groups were predominantly male, whereas high fentanyl intake groups were predominantly female (**Suppl. Fig. 8E**). In contrast, there was no significant association between sex and escalation group membership (p = 0.39) (**Suppl. Fig. 8F**). Further detail on the examination of sex and latent growth curve modeling is provided in **Suppl. File 1**.

### Relationship between acquisition and escalation groups

Because group-based trajectory models of acquisition and escalation phases were constructed independently, we examined the association between acquisition and escalation group assignments. A Chi-Square test for association between acquisition and escalation group assignments yielded p=0.14. Therefore, relatively lower or higher number of presses in the escalation phase was <u>not</u> significantly associated with lower or higher presses during acquisition, highlighting a dissociation between behavioral measures of fentanyl seeking under short and extended access conditions.

### Differential Gene Expression among Escalation Groups

Bulk RNA sequencing identified 17337 unique genes across the three escalation groups (N=8 animals/group) established by the group-based trajectory modeling. Genes with 10 or fewer total counts across all 24 samples were filtered out for pyDESEQ analysis, leaving 14277 identified genes. We examined differential gene expression in high vs low (group 3 vs 1), high vs medium (group 3 vs 2), and medium vs low escalators (group 2 vs 1). Traditional sequencing analysis approaches indicated that 10 genes had significantly increased expression (FDR < 0.05 and log2(FC) > 0.5) in Group 3 compared to Group 1 rats (**Fig. 3A**), including genes coding for voltage-gated potassium and NMDA receptor channels. An additional 37 genes were identified in Group 3 vs Group 1 with an FDR < 0.05 and |log2(FC)| ≤ 0.5 (32 upregulated and 5 genes down regulated). A single gene, *Fgf1* (fibroblast growth factor1), was identified with statistically significant differential expression in Group 3 vs Group 2, (**Fig. 3B**), and no genes were differentially expressed in Group 2 vs Group 1 at a statistically significant level (**Fig. 3C**). Differential expression data for all identified genes in the three comparisons can be found in **Suppl. File 2**.

**Figure 3.**
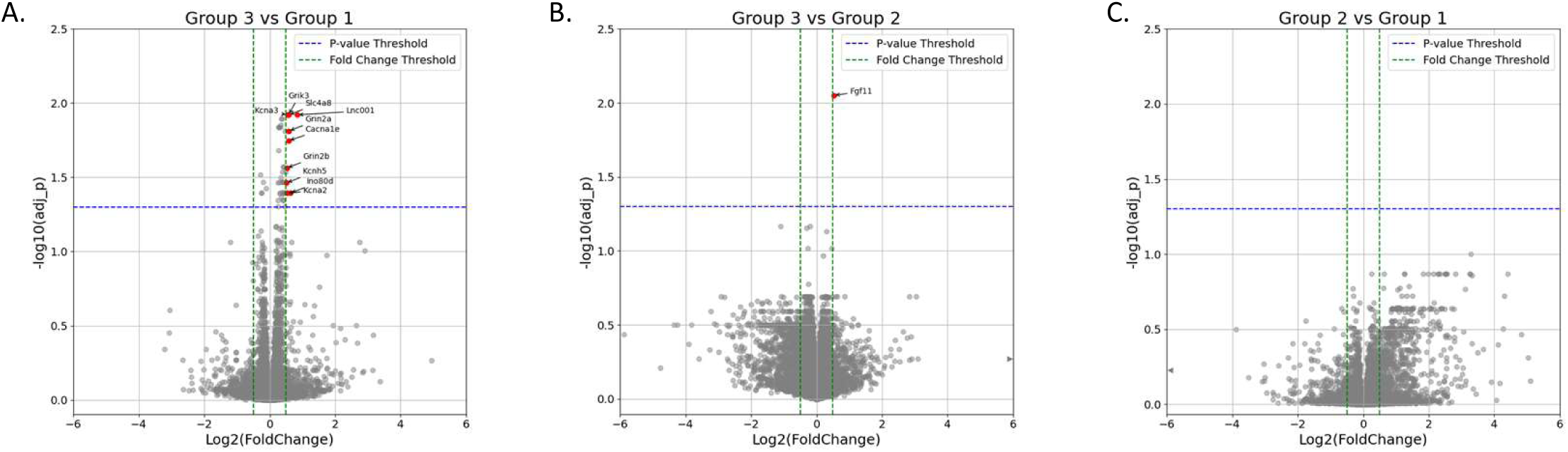
Differential Gene Expression in Fentanyl Escalating Rats. (**A**) Volcano plot of DEGs in fentanyl escalation group 3 vs group 1. (**B**) Volcano plot of DEGs in fentanyl escalation group 3 vs group 2. (**C**) Volcano plot of DEGs in fentanyl escalation group 2 vs group 1. Adjusted p values in volcano plots were calculated according to the pyDESeq2 pipeline. Significantly upregulated genes (log2fold change > 0.5 and FDR<0.05) are colored in red. Blue dotted lines are significance thresholds (-log10(0.05)) and green lines are cut-offs for the log2fold change (log2fold change =0.5 and-0.5).

When using permutations to calculate unadjusted p-values, no statistically significant results remained for individual genes after adjustment for multiple comparisons (**Suppl. Table 1)**. In comparing the three main escalation groups, there were 1056 genes (7.4%) with unadjusted p-values less than 0.05, suggesting that differential expression may exist (as 7.4% is greater than 5%). However, the smallest adjusted p-value for any individual gene was 0.36. In pairwise comparisons between escalation groups, adjusted p-values for all genes were at least 0.19. When looking at the raw numbers of fentanyl infusions during the last three days as a numeric measure of behavior (instead of groups derived from trajectory analysis), one gene had an adjusted p-value of 0.071, while all other genes had adjusted p-values of at least 0.271.

### Identification of Gene Modules Associated with High Fentanyl Escalation

We then performed a signed WGCNA to take into account directionality of the correlation in the gene co-expression network, and three modules (Whitesmoke – 282 genes, Brown – 335 genes), and Lightgrey – 144 genes) were significantly correlated to fentanyl escalation group with no modules correlated to sex (**Fig. 4A,B**). Specifically, all three modules were positively correlated with Group 3 assignment [Whitesmoke (R = 0.5522, p = 0.005), Brown (R = 0.5418, p = 0.006), and Lightgrey (R = 0.4837, p = 0.017)]. The Brown module was negatively correlated with Group 1 assignment [R =-0.5361, p = 0.007], while the negative correlations of Whitesmoke (R =-0.3884, p = 0.061) and Lightgrey (R =-0.3378, p = 0.106) with Group 1 assignment did not reach statistical significance (**Fig. 4B**). The Brown module included 6 of the 10 differentially expressed genes with FDR < 0.05 and log2(FC) > 0.5 and 13 of 37 differentially expressed genes with FDR < 0.05 and |log2(FC)| < 0.5 in the Group 3 rats compared to Group 1. The Whitesmoke included 2 of the 10 differentially expressed genes with FDR < 0.05 and log2(FC) > 0.5 and 14 of 37 differentially expressed genes with FDR < 0.05 and |log2(FC)| < 0.5 in the Group 3 rats compared to Group 1. The Lightgrey module contained no genes with FDR < 0.05 in the Group 3 rats compared to Group 1. Gene lists for each module can be found in **Suppl. File 3,** and heat maps showing correlation between all modules and each sample can be found in **Suppl. Fig. 11**.

**Figure 4.**
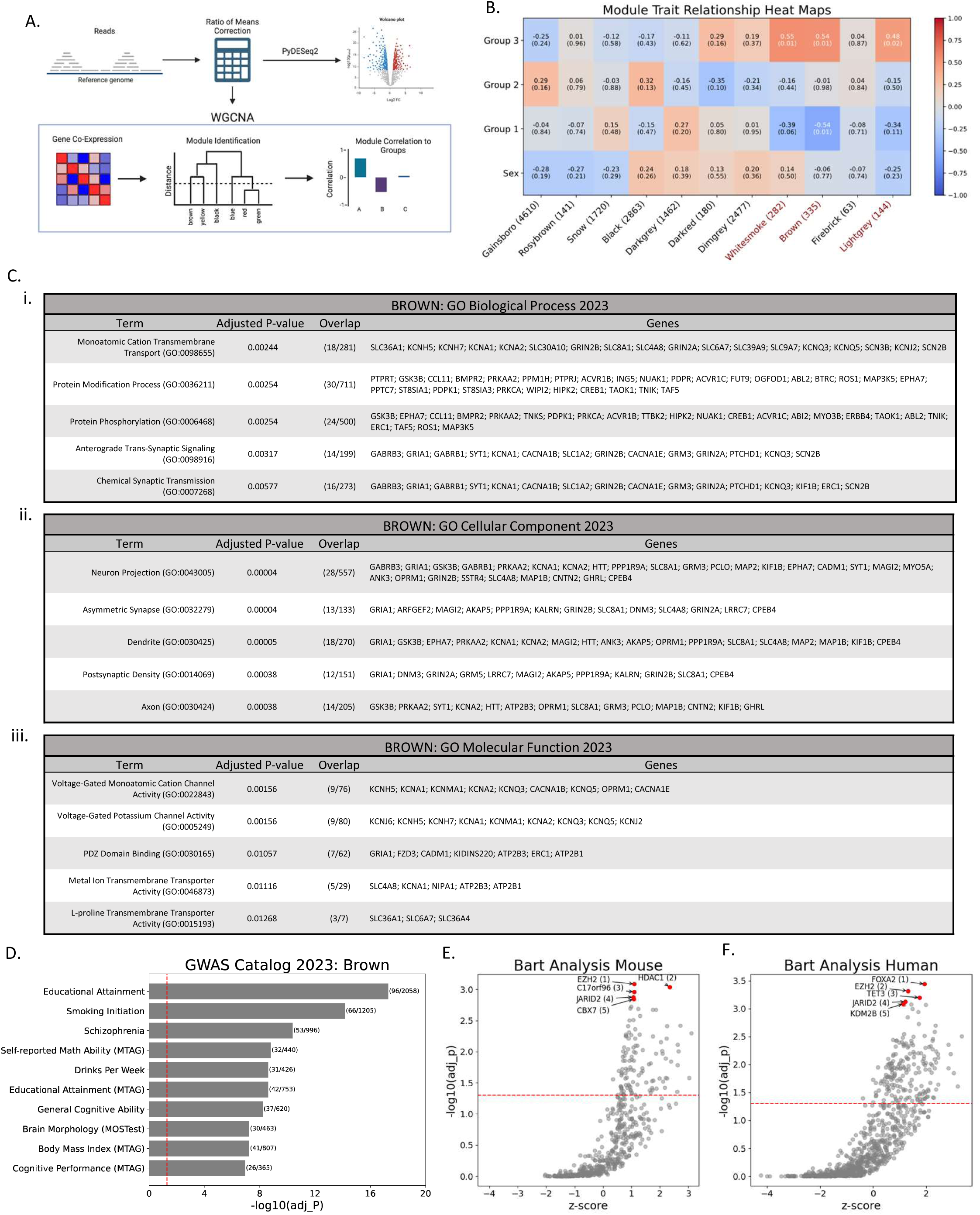
Identification of Gene Modules Associated with High Fentanyl Escalation. (**A**) Graphical representation of the approach of pyDESeq2 and PyWGCNA. (**B**) Heatmap displaying the module-trait relationship for each eigengene, color-coded for directionality of expression (downregulated – blue, upregulated – red). The first three rows indicate group-based model assignments of fentanyl escalation phenotypes. The top value in each box is R and the values in parentheses are P values, with significant values (P<0.05) displayed in white. (**C**) Function enrichment analysis result tables of Brown module conducted via enrichr, (**C.i**) GO Biological Processes 2023, (**C.ii**) GO Cellular Component 2023, (**C.iii**) GO Molecular Function 2023. Term columns are the GO terms, Adjusted P-value are the corrected p-values, overlap is the number of genes in the Brown module that appear in the GO term, and genes is the list of genes from the Brown module in the GO term. (**D**) Function enrichment analysis bar plot of Brown module conducted via enrichr for the GWAS catalog. The y-axis are the GWAS terms, the x-axis is the-log10 of the adjusted p-value, and the parentheses to the right of the bar plot are the overlap of genes from the Brown module and genes in the GWAS catalog term. BART transcriptional regulator predictions generated via the BART mouse library (**E**) and the BART human library (**F**). In **E** and **F** red dots are the top 5 predicted transcriptional regulators.

We next performed functional enrichment analysis of the three modules significantly correlated to fentanyl escalation using the Biological Process, Cellular Component, and Molecular Function ontologies (*29, 30*), as well as the GWAS catalogue (*31*). For the Brown module, there was a total of 46 statistically significant terms across all three ontologies (Biological Process, Cellular Component, and Molecular Function). Terms containing potassium channel genes were present in all three ontologies and terms associated with glutamatergic signaling were found in the Biological Process and Cellular Component ontologies (**Fig. 4C**). The Whitesmoke Module had a total of 17 statistically significant terms across all three ontologies and included potassium channel genes, but not glutamatergic signaling genes (**Suppl. Files 4**). In the Lightgrey module, a single Biological Process term (Lipid Oxidation (GO:0034440)) was significant, and no significant terms were identified in the Cellular Component or Molecular Function ontologies (**Suppl. Files 4**).

Finally, we explored if any of the modules were enriched for genes that have SNPs associated with SUDs via the GWAS catalog accessed via the GSEAPY enrichr function. The Brown module had 157 GWAS catalogue terms that were statistically significant, and 3 of the top 5 GWAS catalog terms were strongly associated with substance use disorders: smoking initiation, schizophrenia, and drinks per week (**Figure 4D**). While the total number of statistically significant GWAS catalog terms for the Brown Module is rather large, this was not the case for all modules, as the Whitesmoke module only had 10 GWAS catalogue terms that were statistically significant, the top term being schizophrenia (**Suppl. Files 4**). The Lightgrey module had 5 significant GWAS catalogue terms that were mostly related to red blood cell characteristics (**Suppl. Files 4**). Furthermore, of the 8 modules identified via the WGCNA that were <u>not</u> correlated to fentanyl escalation group, only one (Black) had any statistically significant GWAS catalogue terms (49), many of which were associated with immune function (**Suppl. Files 4**).

### Transcriptional drivers of fentanyl escalation

To further explore potential mechanisms driving phenotypic differences in fentanyl escalation, we used Binding Analysis for Regulation of Transcription (BART) to predict transcription factors regulating each module based on published mouse and human chromatin immunoprecipitation-sequencing (ChIP-Seq) libraries (*32*), since rat libraries are not yet available. For the Brown module, two of the top 5 predicted transcription factors overlapped in mouse and human libraries: enhancer of zeste 2 polycomb repressive complex 2 subunit (EZH2) [mouse: absolute rank = 1, p < 0.001; human: absolute rank = 2, p < 0.001] and jumonji and AT-rich interaction domain containing 2 (JARID2) [mouse: absolute rank = 4, p = 0.001; human: absolute rank = 4, p < 0.001] (**Figure 4.E,F**). EZH2 is a core subunit of the Polycomb repressive complex 2 (PRC2) and is prominently involved in cancer (*38, 39*). Consistently, in the Whitesmoke module, EZH2 was also in the top 5 predicted transcription factors [mouse: absolute rank = 5, p = 0.002; human: absolute rank = 5, p < 0.001], while JARID2 was in the top 15 [mouse: absolute rank = 14, p = 0.001; human: absolute rank = 11, p = 0.003] (**Suppl. Files 5**). Neither transcription factor reached significance in the Lightgrey module in results from the human or mouse libraries.

### Modeling Contributions of Escalation Risk Genes in Biological Replicates

Finally, we conducted RT-PCR on a select number of genes identified in the Brown module in biological replicates of rats from Groups 1, 2, and 3. Given that the Brown module was positively correlated with Group 3 and negatively correlated with Group 1, we combined the differential expression data of Group 3 versus Group 1 with the Brown module to generate a priority scores for each gene within the Brown module (**Suppl. File 6**). We selected 4 potassium channels (*kcna2, kcnma1, kcnh7, kcnq3*) in the top 25 genes given that function enrichment analysis of the Brown module highlighted terms associated with potassium channel activity and prior evidence implicating potassium channel gene dysregulation in substance use disorders (*40*). We additionally investigated 3 genes in the top 25 associated with glutamatergic signaling (*grin2a*, *grin2b*, *slc1a2*) given the highlighted terms associated with glutamatergic signaling, and 2 genes in the top 25 associated with GABAergic interneurons (*erbb4*, *tnr*) (**Fig. 5.A-I**).

**Figure 5.**
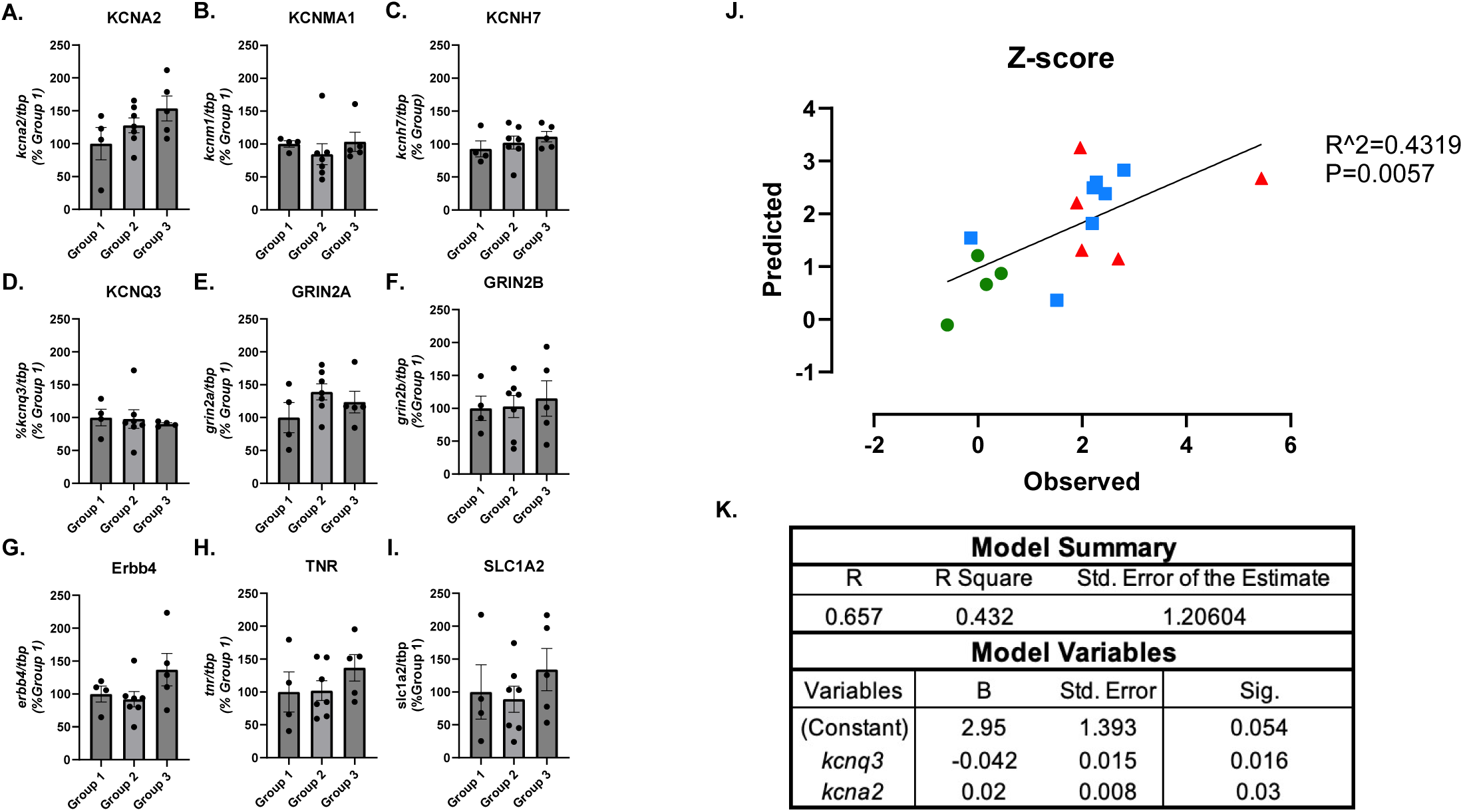
**Testing of Escalation Risk Genes in Biological Replicates**. (**A-I**) Bar graphs show qPCR analysis of select potassium channels, glutamatergic signaling, and GABAergic interneuron associated genes in the three escalation groups. (**J**) Linear regression of predicted versus observed total z-score value. Predicted values were generated using gene expression values and combined lever press z-score calculated from the last 7 days of self-administration. (**K**) Coefficients used in the gene expression model to predict the combined z-score values. *Kcna2*, Potassium Voltage-Gated Channel Subfamily A Member 2; *Kcnq3*, Potassium Voltage-Gated Channel Subfamily Q Member 3. [n = 4 to 7 per group; Figures 5A-I are mean±SEM].

Similar to the conclusions of permutations-based DEG analyses, no individual gene in the PCR data reached statistical significance in a one-way ANOVA treating fentanyl escalation group as the factor. However, linear regression analyses found that expression levels of *kcnq3* and *kcna2* predicted the combined lever press *Z-score* (R^2^ = 0.432; F(2,13) = 4.942, p = 0.025, see Methods). The predicted z-score values from the last seven days of self-administration significantly correlated to the observed values (R^2^ = 0.4319, F(1,14) = 10.64, p = 0.006) (**Fig. 5.J, K)**. Similar to the z-score approach, linear regression analyses using qPCR data and PC1 of each rat also found expression of *kcnq3* and *kcna2* having statistically significant predictive validity of respective PC1 values (R^2^ = 0.435; F(2,13) = 5.014, p = 0.024; **Suppl. Fig. 12**).

## DISCUSSION

A hallmark characteristic of OUD in the DSM-5 emphasizes a user’s overall “loss of control”, which leads to the compulsion to seek and obtain drugs, ultimately resulting in increased consumption(*41*). Preclinical rodent models of escalated drug intake allow rats extended access to fentanyl in SA sessions, resulting in heterogeneity of intake across rats, and individual differences in fentanyl intake may be related to inherent variability in reward sensitivity. First, using latent growth curve models, we found that sucrose breakpoints were not predictive of fentanyl acquisition nor change during the escalation phase, although initial reinforced lever pressing in the escalation phase were predicted by the antecedent sucrose breakpoints. Group-based trajectory modeling was then used to isolate four non-anomalous distinct behavioral phenotypes in the fentanyl acquisition phase and three non-anomalous behavioral phenotypes in the escalation phase. The three-group model following fentanyl escalation was used for transcriptomic analyses for the remainder of the study, and bulk RNA-sequencing of the PFC was utilized to determine differentially expressed genes among the three groups. While a conventional monogenic approach identified a number of genes associated with the endophenotype displaying high escalation of fentanyl intake, the permutation-based monogenic approach revealed no significant associations. Given that complex behavioral responses, such as fentanyl seeking, are likely to develop on the background of cellular plasticity originating from multiple gene loci, a polygenic approach using WGCNA and functional enrichment analysis revealed several genetic targets involving voltage-gated potassium channels, and excitatory glutamate signaling. Further, GWAS catalog functional enrichment results identified modules that were enriched for genes associated with the individual variability to escalate fentanyl. Taken together, identification of gene networks that examine the joint contribution of genes may provide insight into druggable targets that underlie the complex endophenotype associated with the ‘loss of control’ phenomenon seen in OUD.

A major finding in the current study was that breakpoints in the sucrose PR task did not predict subsequent fentanyl self-administration in acquisition sessions or the temporal change in lever presses over the 21-day escalation phase. It is possible that short duration fentanyl sessions limited the ability of sucrose PR to predict fentanyl seeking during the acquisition phase. That is, while each individual rat may possess the phenotypic features that allow for an association between sucrose PR and fentanyl self-administration, a short testing window precluded us from observing such an association. Within opioids, assessments of antecedent sensitivity to non-drug reward have been evaluated using several different paradigms, with mixed results. A study by McNamara, Dalley, Robbins, Everitt and Belin (*42*) was not able to predict escalation in heroin SA with the choice serial reaction time (5CSRT), a task that measures attention and impulsive action in rodents via the ability to withhold a behavioral response to a reward-associated stimulus (*43*). Vasquez, Shah, Re, Laezza and Green (*44*) also found that PR breakpoints for sucrose were not predictive of PR breakpoints for fentanyl SA, although a frustrative non-reward task was able to predict PR breakpoints for fentanyl SA. Contrary to the current findings with fentanyl, unpublished data from our laboratory has shown that antecedent sensitivity to non-drug reward, measured by an identical PR task for sucrose, *did* predict responding for cocaine SA in a PR task (personal communication, Dr. Pavel Ortinski). Similar results showing that discrete traits predict drug intake have been reported when assessing stimulants, as individual differences in baseline impulsive action on the 5CSRT task have predicted greater rates of SA with nicotine (*45*) and cocaine (*46, 47*). Moreover, a study in female rats used a delay discounting task for food reinforcement to screen for high (HiI) or low impulsivity (LoI) and found that HiI rats escalated cocaine intake, while LoI rats did not (*48*). Given the mixed results and pharmacological differences between stimulants and opioids, the inability of non-drug reward sensitivity to predict fentanyl escalation may lend support to the hypothesis that chronic opioid SA *itself* may cause effects on “top-down” control within the PFC (*14*). Moreover, executive control over drug taking becomes progressively weaker as drug experience increases, leading to the “loss of control” phenomenon (*49*). Thus, transcriptomic data may provide insight into the identification of biomarkers that govern this PFC-mediated process of escalation of drug intake.

The failure of a permutation-based monogenic approach to produce candidate genes may have occurred because the ‘loss of control’ phenotype seen in OUD relies on *several* subtle changes within the transcriptomic landscape triggered by escalated fentanyl use. Previous research has demonstrated that diseases such as multiple sclerosis (MS)(*50, 51*), hypertension(*51*), and schizophrenia(*52*) are associated with several genes, and it is the cumulative effects of genes that may result in a summation or interaction of polymorphic changes that drive the disease state. Thus, a polygenic approach using WGCNA was implemented to examine the joint contribution of genes to identify correlation patterns within the three distinct phenotypes ascertained by our model. Within the modules (ME) identified by WGCNA, the correlation pattern within the *Brown* ME is of particular interest. The “addiction-prone” high escalators in Group 3 were positively correlated to the *Brown ME* (0.54), while the low escalating Group 1 rats were negatively correlated (-0.54), and Group 2 showed virtually no directionality (-0.01). A functional enrichment analysis then identified several gene ontologies, including Voltage-Gated Monoatomic Cation Channel Activity (GO:0022843) and Voltage-Gated Potassium Channel Activity (GO:0005249), both of which contained genes (*Kcna2, Kcna3, Cacna1e)* that were significantly regulated when comparing the entire transcriptome of Group 3 and Group 1 using *DESeq2*. Interestingly, *Oprm1* was also included in Brown module and is part of the Voltage-Gated Monoatomic Cation Channel Activity ontology, suggesting that activation of the μ-opioid receptors may be a factor in phenotypic high vs low escalator differences. The prominence of voltage-gated potassium channels genes in our analyses is consistent with their central role in regulation of neuronal firing(*53*). Potassium channels have been linked previously to opioids (*54*), as the *Kcnq* (Kv7) activators retigabine and flupirtine, show antinociceptive effects (*55*), as well as enhanced analgesic effects when combined with morphine (*56, 57*). Moreover, whole-cell patch clamp electrophysiology has shown that fentanyl blocks hERG channels in a voltage dependent manner (*58*), an effect also seen with heroin and methadone (*59*). Interestingly, we have previously explored non-opioid effects of potassium channels in high (highS) and low (lowS) sucrose preferring rats. Sequencing data identified 6 differentially expressed potassium channel transcripts in the NAc between highS and lowS rats, and pharmacological manipulation of the K_V_1.4 current increased responding in a sucrose PR task in lowS rats (*60*).

Transcription factor analysis via BART identified two transcription factors as potential mediators of our transcriptional changes, EZH2 and JARID2. EZH2 is a core subunit of the Polycomb repressive complex 2 (PRC2), which is a complex that regulates methylation of histone H3 lysine 2, with core units of EZH2, EED, SUZ12, RBBP7/4, and acts as a major transcriptional repressor (*38*). Given the integral role of EZH2 in PRC2 histone methylation, it serves multiple important biological functions such as embryonic stem cell differentiation (*61–63*), nervous system development (*64, 65*), immune cell development (*66*), skeletal formation (*67*), and axon regeneration in the peripheral nervous system (*68*). In adult root ganglia cells, overexpression of EZH2 *in vivo* causes increased axon regeneration following injury and *in vitro* causes suppression of ion channel pathways. Interestingly, overexpression of a modified EZH2 with loss of methylation function, ablating the PRC2’s ability to suppress gene expression via HK27me3, causes an upregulation of ion channels pathways, including three differentially expressed genes that overlap with our Brown module (Kcnma1, Kcnq3, Grin2b)(*68*). This suggests a potential shift from EZH2-mediated histone methylation to a non-methylating role of EZH2 in modulating the observed upregulation of ion channel in group 3 high escalating rats. EZH2 has multiple other non-canonical mechanisms to regulate transcription including complexing with or increasing expression of other transcriptional regulators(*39*) and a previous ATAC-seq study examining the striatum of heroin users found enriched enhancer regions of EZH2(*69*). It is worth mentioning that many of the studies demonstrating co-activation of transcription factors were conducted in cancer cells, as EZH2 is highly implicated in cancer (*39, 70*). The PRC2 can complex with multiple subunits that influence function, such as JARID2 (*71*), which influences embryonic development (*72, 73*) and neurogenesis (*74*), and similar to EZH2, is highly implicated in cancer progression (*75*). JARID2 mutations are associated with neurodevelopmental syndrome (*76*), and SNPs in JARID2 are associated with affective dysfunction during alcohol withdrawal(*77*).

GWAS can provide key insights into interpreting association from a transcriptional data set in a biological and genomic context (*78*). Several significant GWAS terms were generated from the *Brown* module, which was positively correlated with the “addiction prone” high escalating rats in Group 3, that may be associated with SUD such as *Smoking Initiation, Schizophrenia,* and *Drinks Per Week.* Gelernter, Kranzler, Sherva, Koesterer, Almasy, Zhao and Farrer (*40*) conducted a GWAS with 5,697 individuals from the United States with OUD and found the most significant genes were involved in potassium and calcium signaling pathways. Of note, the voltage-gated potassium channel transcripts, *Kcnc1* and *Kcng2,* were significantly associated with single-nucleotide polymorphisms (SNPs). Additionally, *Cacna1c*, a voltage-gated calcium channel, was associated with risk for schizophrenia (*79*) and bipolar affective disorder (*80*), two psychiatric conditions often co-morbid with SUD. Taken together with the transcription factor analyses, this suggests that a narrow group of genes in the *Brown ME* may be driving susceptibility to escalated fentanyl intake. This supposition is further supported by our PCR data. While the permutation-based monogenic approach identified no single gene that was statistically different between the three fentanyl escalation groups, two K^+^ channels, *kcnq3* and *kcna2*, were found to predict the combined lever press Z-score over the last 7 self-administration sessions.

In conclusion, we sought to determine whether non-drug reward sensitivity could predict fentanyl escalation, and to determine the transcriptomic signatures within the PFC that are associated with the “loss of control” phenomenon seen in OUD. The gene networks associated with fentanyl escalation highlighted in the present study may represent concordant genomic changes essential to the “loss of control” endophenotype present in OUD. Furthermore, exploratory studies assessing the potential of these gene expression changes as drug discovery targets for OUD, namely CNS excitation networks via voltage-gated potassium channels may lead to new classes of therapeutics for substance use disorders.

## ACKNOWLEDGEMENTS/DISCLOSURES

This work is supported by: NIH R01 DA053070; U01 DA051377; F31DA057812; the University of Kentucky SUPRA Faculty Pilot Grant; T32 DA035200; AFPE regional award; the University of Kentucky SUPRA Graduate/Professional Student Pilot Grant; AFPE Pre-Doctoral Fellowship.

All authors report no biomedical financial interests or potential conflicts of interest.

All data needed to evaluate the conclusions in the paper are present in the paper and/or the Supplementary Materials.

