## Supplementary material for "Behavioral and genetic markers of susceptibility to escalate fentanyl intake": Suppl File 1

**Supplementary Methods**

*Latent growth curve modeling (21-day “6H” experiment*).

We tested a null hypothesis that *z-scores* could be omitted from the model; if this null hypothesis were rejected, we would then test two subsidiary null hypotheses, one asserting that *z-scores* did not directly affect initial positions of latent growth curves and one asserting that they did not directly affect temporal changes. Next, we altered the latent growth curve model to allow coefficients to depend on animal gender. We re-tested the null hypothesis about omitting *z-scores,* and we also tested a null hypothesis about omitting gender; again, rejection of a null hypothesis would trigger two subsidiary tests. A final version of the latent growth curve model permitted interactions between *z-scores,* and sex.

As post-hoc analyses, we pursued latent growth curve modeling for futile presses and for inactive presses, in both 7-day acquisition and 21-day escalation phases. For consistency with our group-based trajectory modeling, futile presses and inactive presses were log transformed for analyses of data from the 21-day escalation phase.

Part of the data exploration was to confirm that we did not need the intercept of the latent growth curve model to include a random effect for the squad to which an animal belonged or for left or right placement of the active lever.

In mathematical formulation, the expected number of fentanyl infusions at time t (measured in days after the first session, so t = 0 at first session, t = 1 at second session, etc.) for an animal with sucrose preference Z score = z was (a + bz) + (c + dz) t + (e + fz) t^2. So, for instance, an animal at the mean on sucrose preference (Z = 0) was expected to follow the pattern a + c t + e t^2, while an animal one standard deviation above the mean (Z = 1) was expected to follow the pattern (a + b) + (c + d) t + (e + f) t^2. The test of no direct dependence of coefficients on Z scores had null hypothesis b = d = f = 0. The two subsidiary null hypotheses were b = 0 (pertaining to initial positions) and d = f = 0 (pertaining to temporal changes).

The altered latent growth curve model expressed the expected number of fentanyl infusions for an animal with sucrose preference Z score = z and sex x (1 for female, 0 for male) as (a + bz + gx) + (c + dz + hx) t + (e + fz + ix) t^2. The test of no direct dependence of coefficients on Z scores again had null hypothesis b = d = f = 0, but in the altered model the test was interpreted as being adjusted for sex; the relevant test for sex had null hypothesis g = h = i = 0, with the interpretation of being adjusted for sucrose preference.

The final latent growth curve model had the structure (a + bz + gx + jzx) + (c + dz + hx + kzx) t + (e + fz + ix + lzx) t^2. This simplifies to (a + bz) + (c + dz) t + (e + fz) t^2 for male animals and to ({a + g} + {b + j} z) + ( {c + h} + {d + k} z ) t + ( {e +i} + {f + l} z ) t^2 for female animals. The parameters j, k, l allowed the association of sucrose preference with fentanyl infusions to differ by sex, and j = k = l = 0 represented a null hypothesis of no interaction between sucrose preference and sex.

When using SAS PROC MIXED to fit a latent growth curve model for inactive presses from the 7-day acquisition phase, we encountered a warning that the estimated variance/covariance matrix of random effects was not positive definite. Therefore, unlike the other latent growth curve models for which results appear in the present article, the latent growth curve models for inactive presses from the 7-day acquisition phase were simplified to include only random intercepts; random adjustments to linear and quadratic coefficients were eliminated.

*Group-based trajectory modeling (21-day “6H” experiment)*.

Provisionally assuming that all trajectories were quadratic, we found that a model with 4 groups exhibited a much better log likelihood than a model with 3 or fewer groups. We then ascertained whether quadratic trajectories could be simplified to linear or even horizontal trajectories, using a backward elimination procedure. Four of the quadratic trajectories were reduced to linear and one was reduced to horizontal. Next, we re-ran the group-based trajectory model with the additional feature that sucrose preference could directly influence probabilities of group membership; a likelihood ratio test was performed for sucrose preference . Finally, we re-ran the model with sex, rather than sucrose preference, being permitted to influence probabilities of group membership.

Examining the three outcomes simultaneously prevented splintering of the sample into excessively many phenotypes. For instance, if we had examined outcomes individually and discovered 3 groups for each outcome, then there would have been 3 x 3 x 3 = 27 combinations of groups, which would have been difficult to interpret. Examining the three outcomes simultaneously, as we did, may have resulted in some phenotypes being unrecognized; however, the phenotypes that were captured presumably described, at least approximately, the vast majority of animals.

We considered two possibilities besides log transforming futile presses and inactive presses, namely: (i) to tolerate the non-normality and proceed without log transformations; and, (ii) to use Poisson rather than censored normal model specifications for futile presses and inactive presses. However, attempts at both possibilities yielded error messages that the likelihood could not be computed.

We were unable to incorporate random effects into the group-based trajectory modeling, so observations for a given animal were regarded as correlated only insofar as they corresponded to the same group; conditional on group membership, observations were regarded as statistically independent.

When re-running the group-based trajectory model to allow sucrose preference (respectively, sex) to directly influence probability of group membership, we supposed that the log probability of being in group k (here k could be 2, 3, or 4) minus the log probability of being in group 1 was linear in sucrose preference (respectively, sex). Thus, the group-based trajectory model was augmented by a generalized logit model with 3 (i.e., one less than the number of groups) slope coefficients; this is why the chi-square distribution on 3 degrees of freedom was used to obtain the critical value for the change in twice the log likelihood.

*Permutation-based testing for single gene differential expression*

We used permutations to obtain unadjusted p-values for differential expression of genes. When comparing two or three escalation groups to each other, we used the following steps to obtain unadjusted p-values. First, we excluded any gene for which adjusted counts within one or more of the three main escalation groups were all zeroes; this left us with 14222 genes. Second, we used the delta method to estimate variances and covariances of log fold changes; direct log transforms of adjusted counts themselves were not usable because some adjusted counts were zeroes. Third, we used the aforementioned estimated variances and covariances to construct a test statistic for differential expression; this was structured like a Hotelling T^2 statistic, but (with non-normally distributed adjusted counts and small samples) we did not assume that the test statistic could be calibrated to a known distribution. Fourth, in each of 10,000 repetitions, we calculated the test statistic again, as if adjusted counts had been randomly permuted across animals; the initial unadjusted p-value for differential expression was the proportion of the 10,000 repetitions in which a larger test statistic was obtained than in the third step. Fifth, for any gene with unadjusted p-value less than 0.05 in the fourth step, we re-calculated the unadjusted p-value to greater precision by using 100,000 repetitions. Sixth, for any gene with unadjusted p-value less than 0.001 in the fifth step, we re-calculated the unadjusted p-value to still greater precision by using 1,000,000 repetitions. Adjusted p-values were obtained using the Benjamini-Hochberg (1995) procedure (*1*). More specifically, the adjusted p-value for a particular gene was defined to be the false discovery rate at which we would be ambivalent between retaining and rejecting the null hypothesis of no differential expression for that gene.

When comparing animals to each other based on the numbers of fentanyl infusions during the last three days of the 21-day escalation phase (a post-hoc analysis not based on trajectory groups), we used the following steps to obtain unadjusted p-values. First, we excluded any gene for which adjusted counts were all zeroes; this left us with 14273 genes. Second, we used the lm function in R to calculate a test statistic for differential expression; this was the absolute value of the usual T statistic from linear regression (in this case, regression of adjusted count on number of fentanyl infusions during the last three days), but we did not assume that the test statistic could be calibrated to a T distribution. The last three steps were the same as in the preceding paragraph.

*Identification of Gene Modules Associated with High Fentanyl Escalation*

Differential gene expression via PyDESeq2 performs a within-subject correction to account for differences in read depth via a ratio of means method and across subjects to account for biological variability via estimate of dispersions, and fits generalized linear model to calculate the log2Fold change for each gene(*2, 3*). In contrast, the Weighted Gene Co-Expression Network Analysis (WGCNA) allows the analysis of groups (‘modules’) of genes across experimental conditions by hierarchical clustering of genes identified by the normalized PyDESeq2 counts(*4, 5*). Correlating total gene expression within modules to fentanyl escalation group assignments allowed us to identify modules that are differentially expressed across the three escalation groups

**Supplementary Results**

*Examination of Sex and Latent Growth Curve Modeling*

There were 36 male and 36 female rats that finished sucrose PR training. Among these, the fentanyl acquisition phase was completed by 32 male and 31 female rats, while the escalation phase was completed by 28 male and 30 female rats.

As for potential sex differences, sex was not a factor in association between sucrose PR and reinforced lever pressing in latent growth-curve models in the analyses above, as no interaction was detected between sucrose preference and sex in either the acquisition (p=0.629) or escalation phase (p=0.20). However, sex was found to predict reinforced lever pressing during the acquisition phase (p=0.02) in a model that adjusted for sucrose preference. More specifically, sex predicted the change over time in the number of infusions (p=0.043), but not the starting number of infusions (p=0.255, **Figure 5A-B**). During the escalation phase, sex was also found to predict reinforced lever pressing for fentanyl (p=0.006). In this case, sex predicted both the initial number of infusions (p=0.045) and the change over time in the number of infusions (p=0.006, **Figure 5C-D**). Latent growth curve models examining inactive and futile lever responding as a function of sex and sucrose PR can be found below.

In latent growth curve models, sucrose PR performance did not predict inactive lever pressing for fentanyl without (p = 0.929) and with (p = 0.928) adjustment for sex. However, sex was significantly associated with inactive pressing in acquisition (p = 0.012), both in terms of initial position (p = 0.001) and temporal change (p = 0.023). Moreover, a highly significant sucrose/sex interaction was detected (p < 0.001), relating to both initial position (p < 0.001) and temporal change (p < 0.001) (**Suppl. Fig.9A-B**). The nature of the interaction was that, among females, sucrose seeking was significantly associated with inactive pressing (p = 0.004), concerning both initial position (p = 0.001) and temporal change (p = 0.012). Surprisingly, females with low sucrose seeking began with more inactive presses but declined rapidly; this finding was driven largely by three females whose first-day inactive presses were at least three times the mean among all females. Among males, sucrose seeking was not significantly associated with inactive pressing during the acquisition phase (p = 0.097).

During the escalation phase, sucrose lacked statistical significance without (p = 0.610) and with (p = 0.678) adjustment for sex, sex was not significantly associated with inactive pressing (p = 0.570), and no significant sucrose/sex interaction was detected (p = 0.070) (**Suppl. Fig. 9C-D**).

In latent growth curve models of futile pressing, sucrose lacked statistical significance without (p = 0.835 acquisition; p = 0.534 escalation) and with (p = 0.811; p = 0.665) adjustment for Sex. Sex was significantly associated with futile pressing (p < 0.001; p < 0.001), both in terms of initial position (p = 0.029; p < 0.001) and temporal change (p = 0.032; p < 0.001). Significant sucrose/sex interactions were not detected (p = 0.728; p = 0.424). Essentially, females started higher on futile presses and rose more quickly during acquisition, while males then rose more quickly during escalation (**Suppl Fig.10A-D**).

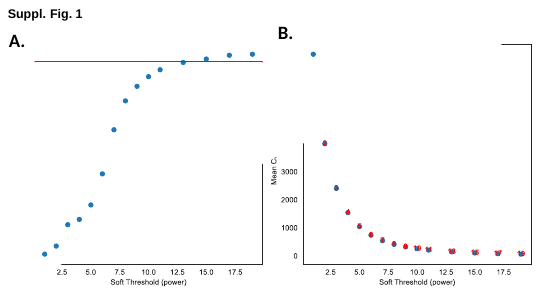
Fig. S1. Evaluation of soft threshold value for scale free network.

(**A**) Graphical representation of correlation between log(k) and log(p(k)), where k is number direct connections of a given gene to all other genes, and p(k) is the frequency distribution of connectivity. The y-axis is the correlation between log(k) and log(p(k)) for a given soft threshold value, which is displayed on the x-axis. The red bar is R^2^ = 0.9, and soft threshold value was selected where curve plateaus above R^2^ value of 0.9. (**B**) The average connectivity of all genes, y-axis, inputted into WGNA function under increasing soft threshold values, x-axis.

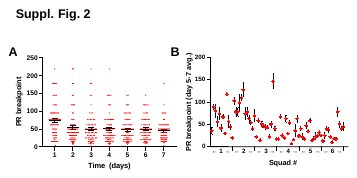

Fig. S2. Individual data points of sucrose progressive ratio self-administration.

(**A**) Average sucrose breakpoints across the 7 progressive ratio sucrose self-administration sessions. (**B**) Average breakpoint for each rat over the last 3 progressive ratio sucrose self-administration session broken down by squad, with each squad containing 12 rats. [n=72, error bars indicate SEM]

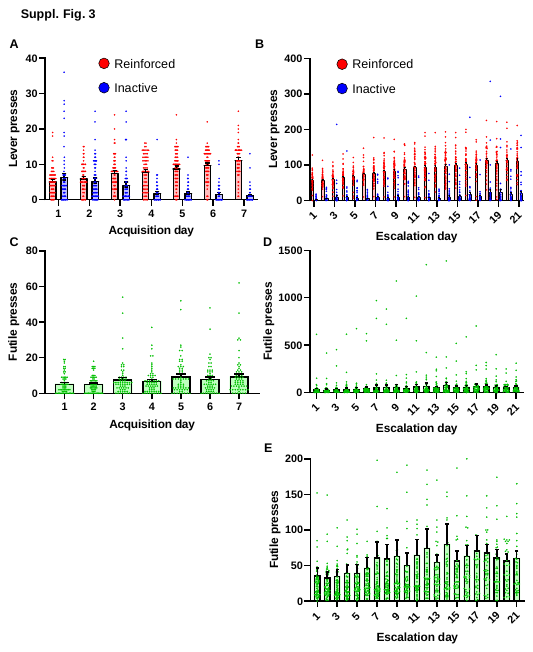

Fig. S3. Individual data points of fentanyl self-administration.

(**A**) Average reinforced and inactive lever presses during the acquisition sessions. (**B**) Average reinforced and  inactive lever presses during escalation sessions. One value of 4042 inactive presses on day 2 is excluded. (**C**) Average futile lever presses during the fentanyl acquisition sessions. (**D**) Average futile lever presses during the fentanyl escalation sessions. (**E**) Data identical to D, with truncated Y-axis to highlight the means. [n=63, dots indicate individual rats and error bars indicate SEM]

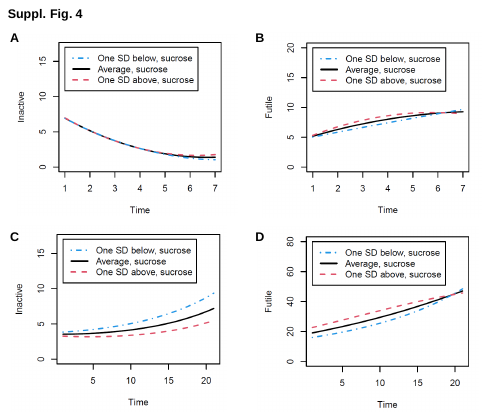

Fig. S4. A modeled relationship between sucrose PR and inactive/futile lever presses during fentanyl acquisition and escalation.

Latent growth curve model considers hypothetical animals behaving exactly at the average breakpoint over the last three days of sucrose PR, one standard deviation above the average, and one standard deviation below the average. The relationship between sucrose PR and lever presses is visually appreciated as the magnitude of difference between the 3 growth curves, corresponding to hypothetical animal’s average (black), one SD above (red) and one SD below (blue) on PR for (**A**) acquisition inactive presses, (**B**) acquisition futile presses, (**C**) escalation inactive presses, and (**B**) escalation futile presses.

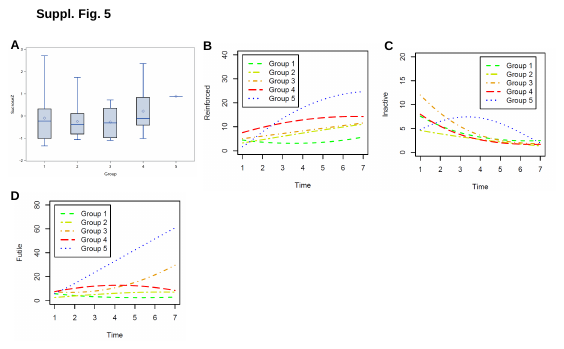

Fig. S5. Five phenotypes of fentanyl acquisition identified by group-based trajectory modeling.

(**A**) Box plot of sucrose breakpoint *z-scores* by assigned acquisition groups. Group assignment for the acquisition phase was based on the following parameters: (**B**) reinforced lever presses, (**C**) inactive lever presses, (**D**) futile lever presses for animals placed in Group 1 (green), Group 2 (yellow), Group 3 (orange), Group 4 (red), and Group 5 (blue).

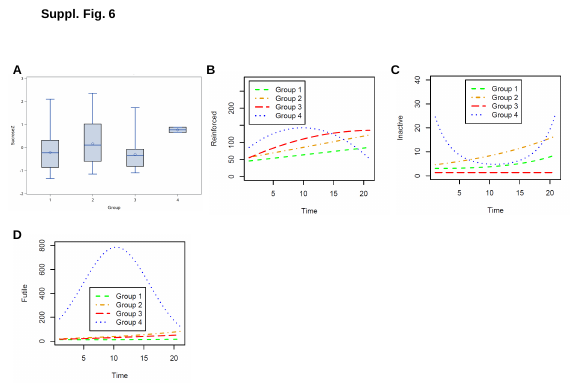

Fig. S6. Four phenotypes of fentanyl escalation identified by group-based trajectory modeling.

(**A**) Box plot of sucrose breakpoint *z-scores* by assigned escalation groups. Group assignment for the escalation phase was based on the following parameters: (**B**) reinforced lever presses, (**C**) inactive lever presses, (**D**) futile lever presses for animals placed in Group 1 (green), Group 2 (yellow), Group 3 (orange), and Group 4 (blue). Error bars in A indicate the range of data.

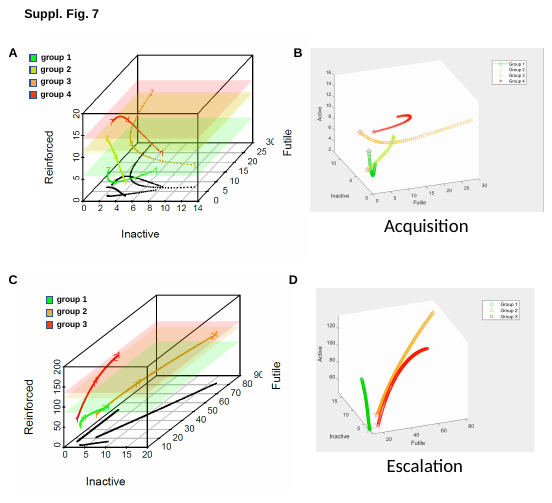

Fig. S7. Three-dimensional representations of acquisition and escalation trajectories.

The temporal direction of lever press changes is represented by numbers interrupting each of the group curves (i.e. days 1-7 and day 4 midpoint for the acquisition groups; days 1-21 and day 11 midpoint for the escalation groups). The 3D length of each curve reflects the extent of change throughout (**A**) acquisition and (**C**) escalation phases. The specific axis displaying these changes can also be appreciated by the ‘shadow’ of the corresponding group (black curves in A and C) shown in a 2D projection of inactive vs futile lever presses. The extent of changes on the reinforced lever press axis can be appreciated by the height of the shaded parallelograms color-coded according to the respective groups. For example, escalation group 3 attains the highest level on reinforced presses while also increasing on futile presses; escalation group 2 attains the next highest level on reinforced presses while also increasing (more than group 3) on futile presses and on inactive presses. Video animations are also shown to depict trajectories throughout (**B**) acquisition and (**D**) escalation from different perspectives.

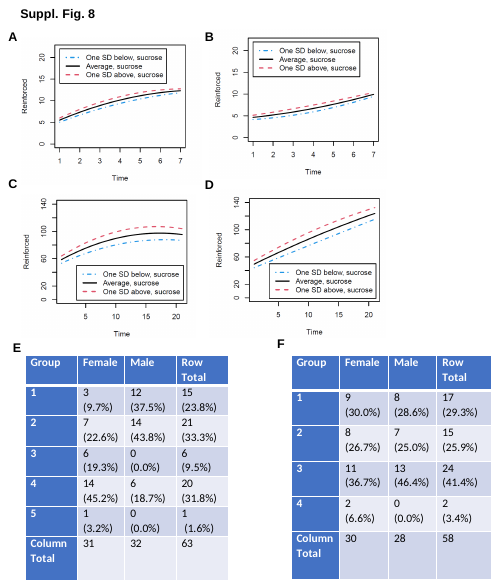

Fig. S8. Latent growth curve modeling of reinforced active lever presses in male and female rats during fentanyl self-administration.

Growth curve models of reinforced presses during fentanyl acquisition by (**A**) female and (**B**) male rats. Growth curve models of reinforced presses during fentanyl escalation by (**C**) female and (**D**) male rats. Animal numbers in acquisition (**E**) and escalation (**F**) models.

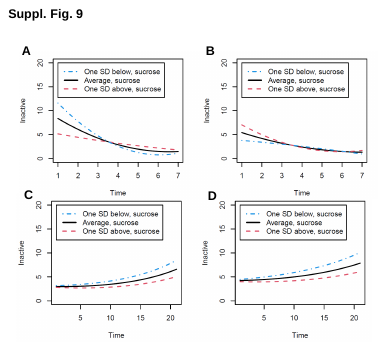

Fig. S9. Latent growth curve modeling of inactive lever presses in male and female rats during fentanyl self-administration.

Growth curve models of inactive presses during fentanyl acquisition by (**A**) female and (**B**) male rats. Growth curve models of inactive presses during fentanyl escalation by (**C**) female and (**D**) male rats.

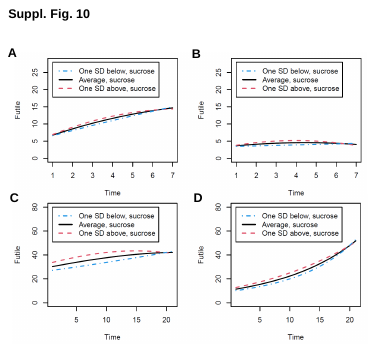

Fig. S10.

**Latent growth curve modeling of futile active presses in male and female rats during fentanyl self-administration.**

Growth curve models of futile presses during fentanyl acquisition by (**A**) female and (**B**) male rats. Growth curve models of futile presses during fentanyl escalation by (**C**) female and (**D**) male rats.

Fig. S11. Module vs sample correlation heatmaps.

(**A-K)** Module expression profiles and heatmaps for each module identified by WGCNA analysis. Columns in each panel represent individual samples. Above, the top two rows represent metadata for each sample (group assignment and sex). The middle row is the bar plot of module expression for each sample. Below is a heat map demonstrating gene expression of each gene in the module for every sample.

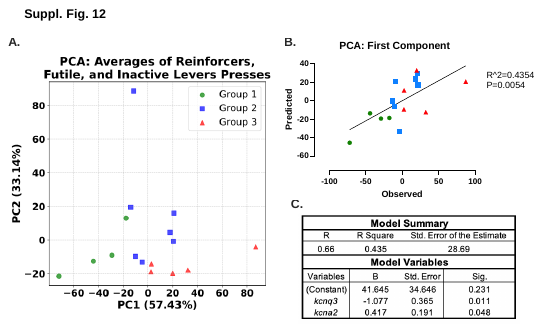

Fig. S12. Principal component analysis for correlation of gene expression and behavioral data.

(**A**) Principal component analysis of the average reinforcers received, inactive lever presses, and futile presses over the last 7 days of self-administration. The x-axis in the first principal component and the y-axis in the second principle component. Values in parenthesis are the amount of variance attributed to each component. (**B**) The graphed predicted versus observed PC1. Predicted values were generated using gene expression values and PC1 calculated from PCA analysis. Statistics presented on graph are the linear regression between predicted and observed PC1. (**C**) The table represents the coefficients used in the gene expression model to predict PC1. *Kcna2*, Potassium Voltage-Gated Channel Subfamily A Member 2; *Kcnq3*, Potassium Voltage-Gated Channel Subfamily Q Member 3. [n = 4 to 7 per group]

| **Comparing gene expression by...** | **Fraction of unadjusted p-values less than 0.05** | **Smallest adjusted p-values**  **and corresponding genes** |
| --- | --- | --- |
| behavioral groups (1 vs 2 vs 3) | 1056/14222 (7.4%) | 0.361 Lnc001; Gja4; Ythdf2; Slc30a10; Mmp8 |
| behavioral groups (2 vs 3) | 1072/14222 (7.5%) | 0.504 Brf1; Fgf11; Ythdf2; Gorasp1; Cuedc1; Gpr107; Gfod2; Nfia; Asic4 |
| behavioral groups (1 vs 3) | 1223/14222 (8.6%) | 0.190 Pik3c2b; Tsc22d2; Gja4; Lnc001; Stk38l; Ddx50 |
| behavioral groups (1 vs 2) | 563/14222 (4.0%) | 1.000 |
| average infusions last three days | 553/14273 (3.9%) | 0.071  RT1-DMa  0.271  Ndufa7 |

Table S1.

Permutation analyses of DEGs.
